# Genomic evidence for supergene control of Darwin’s “complex marriage arrangement” – the tristylous floral polymorphism

**DOI:** 10.1101/2024.01.15.575583

**Authors:** Haoran Xue, Yunchen Gong, Stephen I. Wright, Spencer C. H. Barrett

## Abstract

Tristyly is a polymorphism characterized by three flower morphs with reciprocal stigma and anther heights controlled by two epistatically interacting diallelic loci (*S* and *M*), hypothesized to be supergenes. Chromosome-level genome assemblies of *Eichhornia paniculata* identified the *S*- and *M*-loci. The *S*-locus is a supergene consisting of two divergent alleles: The *S*-allele (2.51Mb) with three *S*-allele specific genes hemizygous in most *S*-morph plants and the *s*-allele (596kb) with five *s*-allele specific genes. Two of the *S*-allele specific genes, LAZY1-S and HRGP-S, were specifically expressed in styles and stamens, respectively, making them tristyly candidate genes. The *M*-locus contained one gene (LAZY1-M), homologous to LAZY1-S, that was present in the *M*-allele but absent from the *m*-allele. Estimates of allele ages are consistent with the prediction that the *S*-locus evolved before the *M*-locus. Re-use of the same gene family highlights the potential role of gene duplication in the evolution of epistatic multilocus polymorphisms.

---

Flowering plants possess a remarkable diversity of floral strategies promoting cross-pollination and outcrossed mating. This variation attracted the attention of Charles Darwin who devoted most of his classic book “*The Different Forms of Flowers on Plants of the Same Species*”^1^ to the floral polymorphism heterostyly in which populations are characterized by two (distyly) or three (tristyly) floral morphs, which possess a reciprocal arrangement of stigma and anther heights^2^. Darwin described heterostyly as a ‘complex marriage arrangement’ promoting cross-pollination between the floral morphs by animal pollinators owing to segregated pollen placement on their bodies (see Fig. 1 in Ref. 3). Contemporary experimental evidence generally supports this hypothesis also finding that heterostyly limits sexual interference between female and male function thus reducing pollen wastage^3–4^. Heterostyly is reported from 28 animal-pollinated angiosperm families^5–6^ and represents a remarkable example of adaptive convergence in form and function. Controlled crosses on over a dozen unrelated species have established that distyly is governed by a single diallelic Mendelian locus (*S*) with dominance, whereas tristyly was controlled by two diallelic epistatically interacting loci (*S, M*), both with dominance^7^ (Fig. 1). To resolve the discrepancy between the complexity of the heterostylous syndrome and its relatively simple inheritance, it has been proposed that heterostyly loci are in fact supergenes comprised of clusters of functionally related genes^8^. Evidence indicates that these genes are tightly linked to each other due to suppression of recombination and are inherited as a single unit^7^.

**Figure 1.**
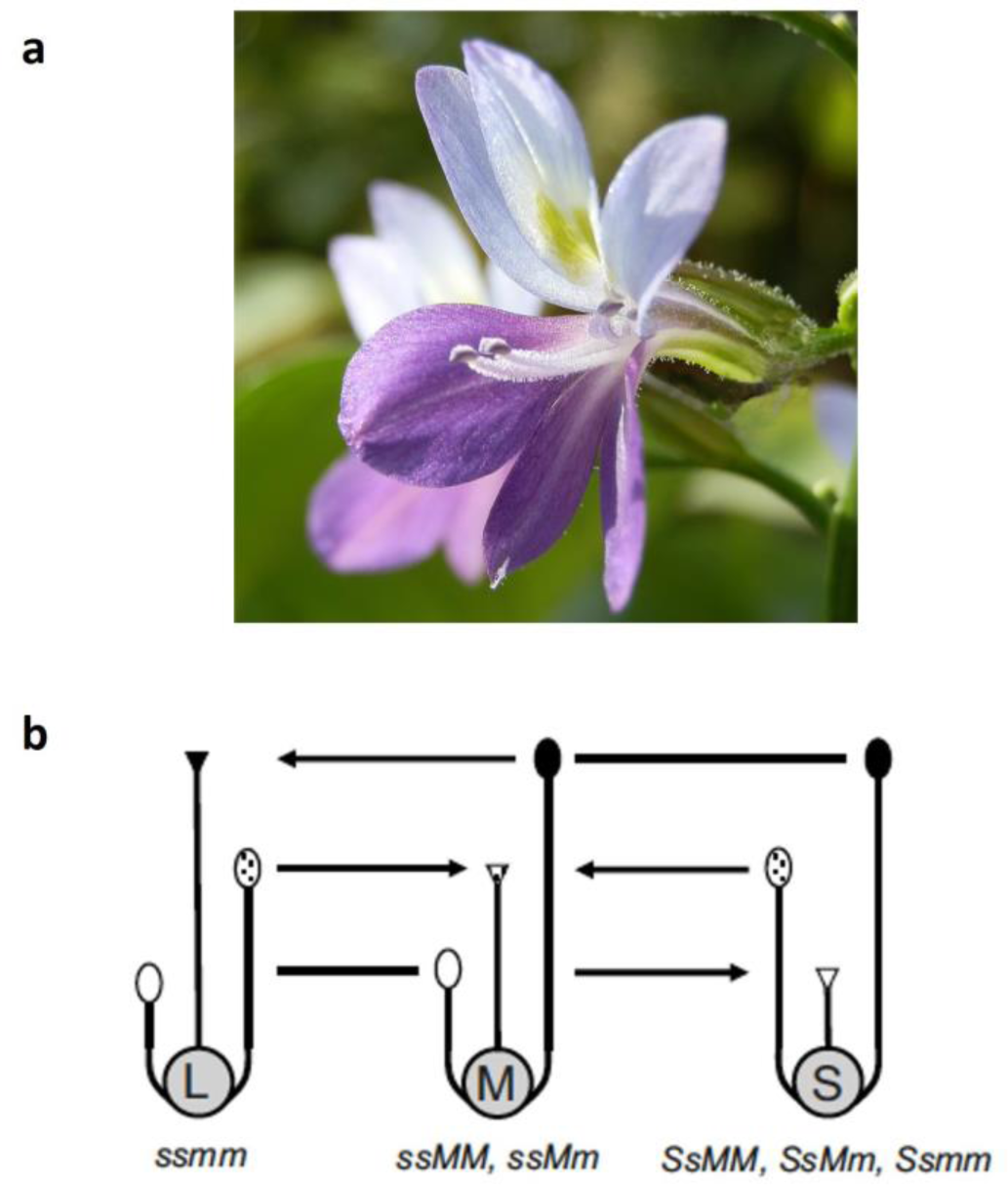
Tristyly in *Eichhornia paniculata*. **a**,Flower of the *S*-morph illustrating exerted long- and mid-level stamens. The short style is concealed in the perianth tube. **b,** The arrangement of styles and stamens in the three floral morphs of a tristylous species. L, M and S refer to the long-, mid- and short-styled morphs, respectively. The arrows indicate “legitimate” cross-pollinations between anthers and stigmas of equivalent height. In most tristylous species these are the only compatible matings because of trimorphic incompatibility. However, in *E. paniculata* “illegitimate” pollinations are permitted as the species is self- and intramorph compatible^35^. Genotypes for the floral morphs of *E. paniculata* are indicated.

Supergenes are also known to control several other complex polymorphic traits, including Batesian mimicry in *Heliconius* butterflies, social organizations of fire ants, and plumage variation and behaviour in ruff^9–11^. Of particular importance for understanding the evolution of supergenes is to determine their genetic architecture and the mechanisms causing suppression of recombination. Identification of the causal loci can also provide important insights into the age of the alleles, to assess the evidence for long-term balanced polymorphisms maintaining variation. Recent efforts have been made to elucidate the genetic architecture of heterostylous supergenes in a handful of distylous systems (reviewed in^12–13^). Significantly, genomic studies on *Fagopyrum, Gelsemium*, *Linum*, *Nymphoides*, *Primula,* and *Turnera*, each belonging to a different angiosperm lineage, have demonstrated that distyly is controlled by several hemizygous genes present in the short-styled morph (*S*-morph) but absent from the long-styled morph (L-morph)^14–20^. For example, in *Primula*, the *S*-locus is a hemizygous region containing five genes controlling the expression of different components of the distylous polymorphism^14–15, 21^.

All molecular investigations of heterostyly have focused on distylous species and the supergene hypothesis has yet to be investigated in any of the six angiosperm families in which the more complex tristyly occurs^22–23^. Charlesworth^24^ suggested that alternatives to the supergene hypothesis should be considered for tristyly because of the absence of empirical evidence for supergenes. Thus, the genetic basis and evolutionary assembly of this two-locus epistatically interacting genetic system remains unclear.

Here, we generate chromosome-level *de novo* genome assemblies with long-read sequences and integrate them with data from population genomic and transcriptomic analysis to investigate the genetic architecture of tristyly in *Eichhornia paniculata*, an annual aquatic native to the Neotropics. Earlier molecular studies demonstrated that the *S*- and *M*-loci in this species are closely linked (separated by a map distance of 2.7 cM), and that the *M*-locus is located on linkage group 5 in a region containing 204 genes^25^, but the genomic location of the *S*-locus was not investigated, and they did not resolve whether the *M*-locus is a supergene or merely contains one pleiotropic gene^12^.

To investigate this issue and provide novel insight into the molecular basis of tristyly we addressed the following questions: 1) What is the genomic architecture of the *S*- and *M*-loci and are they indeed supergenes? 2) What are the likely candidate genes at the two loci controlling different components of tristyly? 3) What is the evolutionary timing of the origins of the *S*- and *M*-loci? We note that in the only theoretical analysis of the evolution of tristyly, the origin of the *S*-locus was inferred to have preceded the origin of the *M*-locus^24^. Our results are consistent with this stepwise origin of the two loci, with recurrent gene duplication from the same ancestral gene playing a key role in the formation of the epistatic interaction between the two loci.

## Results

### Comparisons of chromosome-level genome assemblies

We generated chromosome-level genome assemblies from a heterozygous *S*-morph individual (*SsMm*) and an L-morph individual (*ssmm*) combining long-read PacBio sequencing with proximity ligation (Omni-C and Hi-C) scaffolding (see methods and Fig. S1). In both assemblies, most of the two genomes were assembled into eight major chromosomal scaffolds (Table 1). Comparisons of the two assemblies demonstrated large-scale concordance, with small-scale differences likely reflecting inversions and/or small assembly errors (Fig. S2a). There was clear evidence for a recent whole genome duplication event in which each chromosome showed strong synteny with one or more additional chromosomes. Based on the distribution of synonymous substitution rate (*Ks*) values between paralogs, the most recent whole genome duplication event has a mode of *Ks*=0.38 (Fig. S2b). Assuming a substitution rate of 7 x 10^-9^ per year^26^ and a whole genome duplication event, this would correspond to two sets of homeologous chromosomes that diverged less than roughly 30 million years ago. Since the base chromosome number in Pontederiaceae was inferred to be *n*=8^27^, the duplication event was likely shared across the family, with additional lineage-specific duplications due to additional rounds of polyploidy occurring in other lineages of the family^28^. Unless otherwise noted, our analysis below uses our *SsMm* assembly as the reference genome.

The Marey maps (plots of the linkage map against the physical map) indicated that there was a large (likely pericentromeric) region of low recombination in the middle of each chromosome (Fig. 2a). In chromosomes 3 and 7, the region extended to one end of the chromosome, likely corresponding to submetacentric or acrocentric chromosomes, whereas the remaining chromosomes appeared to be more metacentric. As expected, the regions of reduced recombination were associated with a high repeat density (particularly LTR retrotransposons), low gene density, and low nucleotide diversity, whereas the more highly recombining chromosome tips were gene-dense, comprised of fewer repeats, and has higher nucleotide diversity (Fig. 2b-f, Fig. S3).

**Figure 2.**
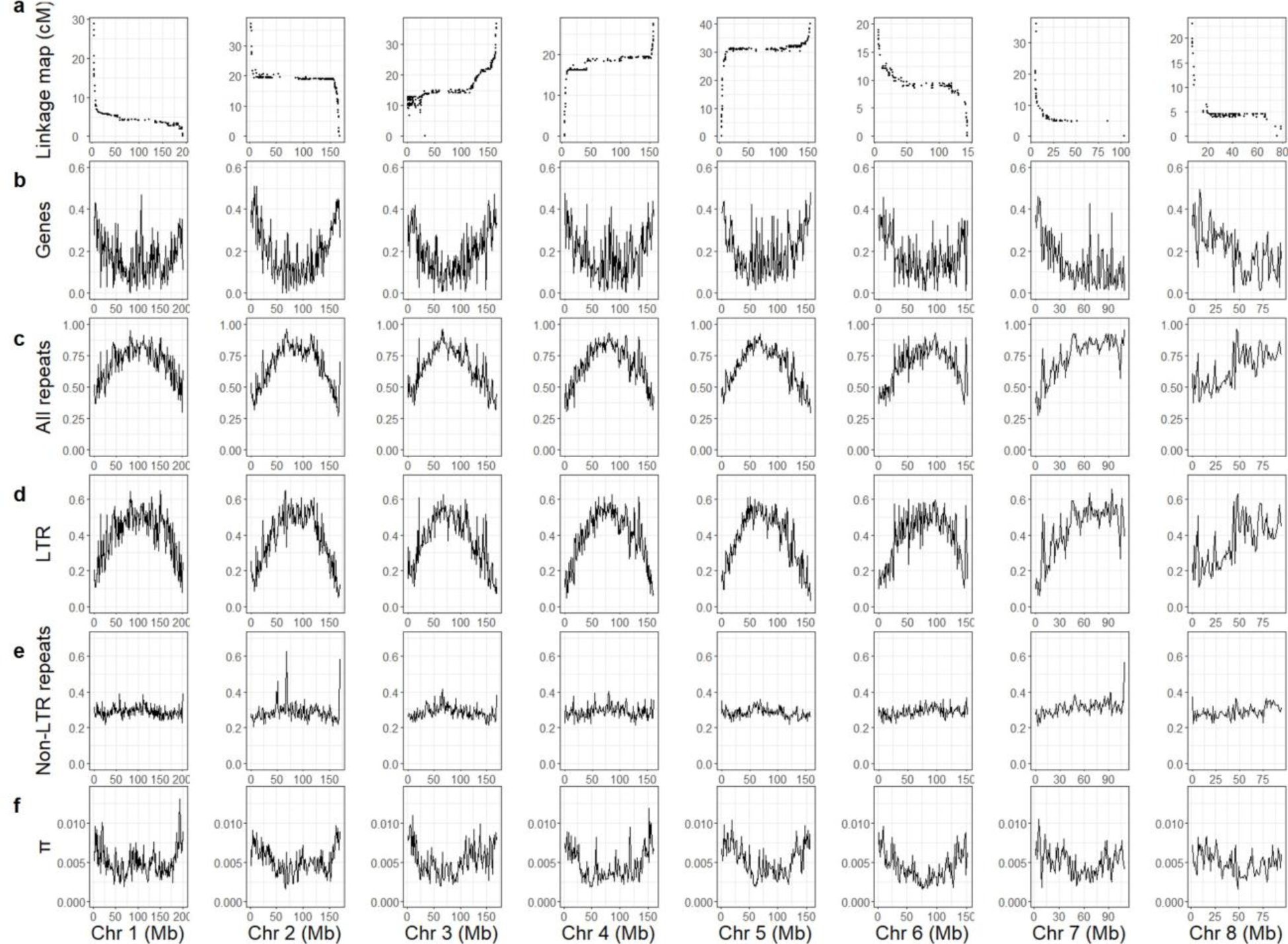
The recombination landscape, genome content, and nucleotide diversity (π) across the chromosomes of *Eichhornia paniculata*. **a**, Marey maps plotting the position on the linkage map (cM) against the position on the physical map (Mb). **b-e,** Densities of genes (**b**), all repetitive elements (**c**), LTR retrotransposons (**d**), and non-LTR repeats (**e**), all measured as proportions of 1-Mb windows. **h,** Nucleotide diversity (π) of 1-Mb windows.

### Both tristyly loci occur in a large low-recombining region

Using a genome-wide association study (GWAS) we identified 4,640 SNPs that were significantly associated with the *S*-morph (*S*---) versus non-*S*-morphs (*ss*--, i.e. L- and *M*- morphs) (*p* < 2.52397779 x 10^-8^, Fig. 3a). 96.8% of these SNPs occurred in a very large 63.1 Mb region on chromosome 2, embedded in an extensive region of reduced recombination on this chromosome (Fig. 3a). As expected from earlier genetic work^25^, the *M*-locus region is partially linked to the *S*-locus. We found 4,541 SNPs significantly associated with the *M*- (*ssM-*) versus L- (*ssmm*) morphs (*p* < 2.23826411 x 10^-8^, Fig. 3b). 98.0% of these SNPs were in a 35.1 Mb region on chromosome 2, at the other end of the region of reduced recombination from the *S*- locus region. Thus, both the *S*- and *M*-loci governing tristyly occur in a large, putatively pericentromeric region of low recombination on chromosome 2 causing extensive and significant SNP associations across many Mb of sequence.

**Figure 3.**
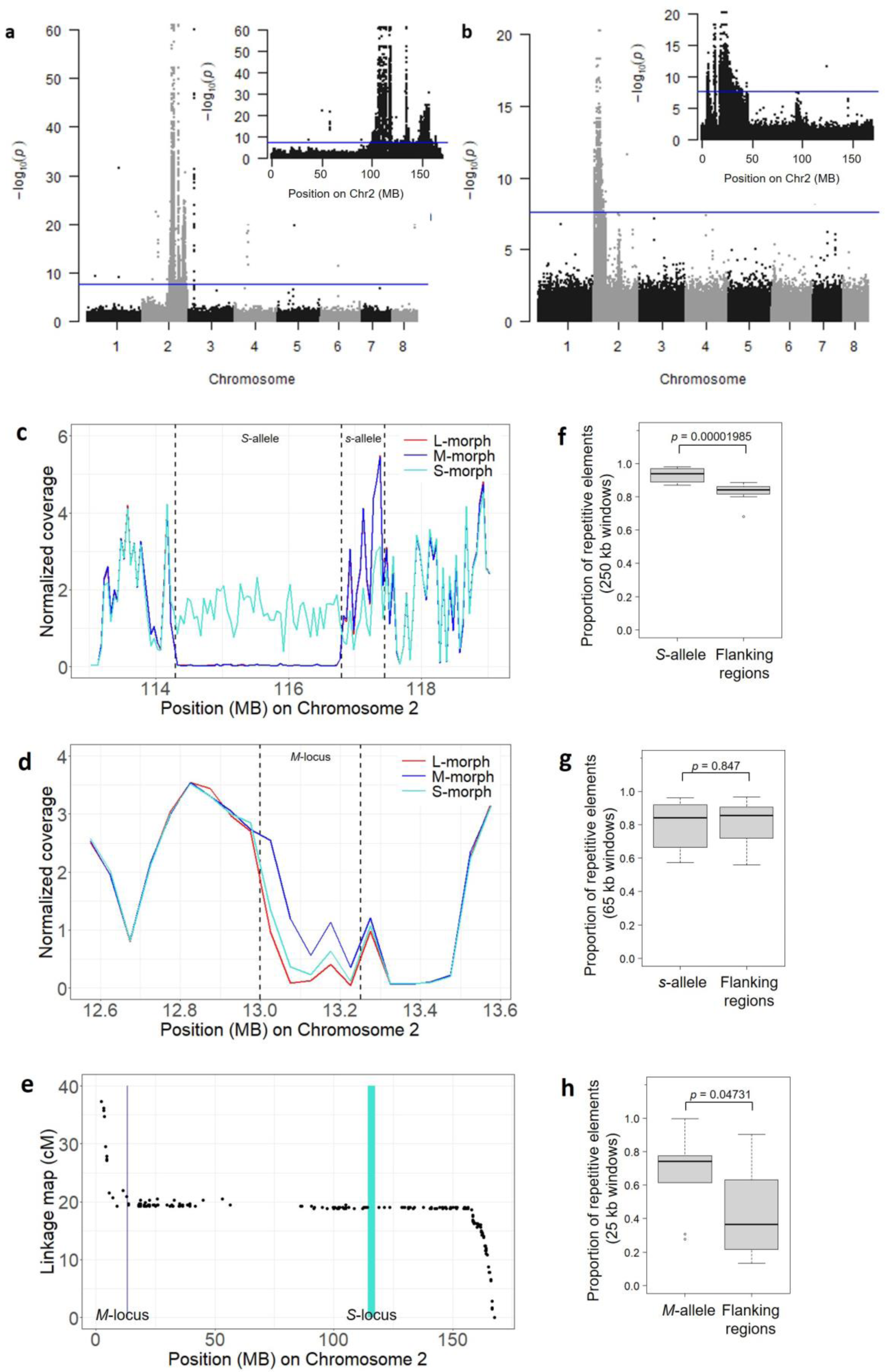
SNPs associated with tristyly in *Eichhornia paniculata* and morph-specific read coverage, recombination landscape, and repeat content of the *S*- and *M*-loci. **a**, SNPs associated with *S*- vs. non-*S*-morphs. The blue lines indicate *p* = 2.52397779 x 10^-8^. Inset panels zoom in on chromosome 2. **b,** SNPs associated with *M*- vs. L-morphs. The blue lines indicate *p* = 2.23826411 x 10^-8^. Inset panels zoom in on chromosome 2. **c**, Average normalized coverages of the L- (red), *M*- (blue), and *S*- (cyan) morph plants at the *S*-locus (*S*- and *s*-alleles assembled next to each other). The data were based on 20 plants per morph. **d,** The average normalized coverages of the L- (red), *M*- (blue), and *S*- (cyan) morph plants at the *M*-locus. The data were based on 20 plants per morph. **e,** The recombination landscape of chromosome 2. **f-h,** Comparisons of proportion of repetitive elements between the *S*-allele (ten 250-kb windows), *s*- allele (ten 65-kb windows), and *M*-allele (ten 25-kb windows), and their flanking regions (five windows upstream and five windows downstream to the locus, with window size equal to the allele being compared with).

### The *S*-locus is a supergene consisting of divergent alleles with hemizygous genes

The *S*- and *M*-locus associated regions we identified were both extremely large with very low rates of recombination. To further narrow down possible candidate genes and to investigate the potential hemizygous regions associated with tristyly, we identified regions with significant differences in read coverage between the floral morphs.

We identified 50 consecutive 50-kb windows that were specific to the *S*-allele (total length: 2.5 Mb; genome coordinates: Chr2:114,300,000-116,800,000), with read coverage significantly higher in 20 *S*-morph (*S*---) plants than 40 non-*S*-morph (*ss*--) plants (criteria: the ratio between the average read coverage of non-*S*-morph and *S*-morph plants was lower than 0.4, with *p*-value of the *t-*test < 9.73217066 ×10^-7^, Fig. 4a). Downstream adjacent to this 2.5 Mb region were 13 consecutive 50-kb windows (total length: 650 kb; genome coordinates: Chr2:116,800,000-117,450,000) that were specific to the *s*-allele and had read coverage significantly lower in 20 *S*-morph (*S*---) plants than 40 non-*S*-morph (*ss*--) plants (criteria: the ratio between the average read coverage of non-*S*-morph and *S*-morph plants was higher than 1.6, with the *p*-value of the *t*-test < 9.73217066 ×10^-7^, Fig. 4a).

**Figure 4.**
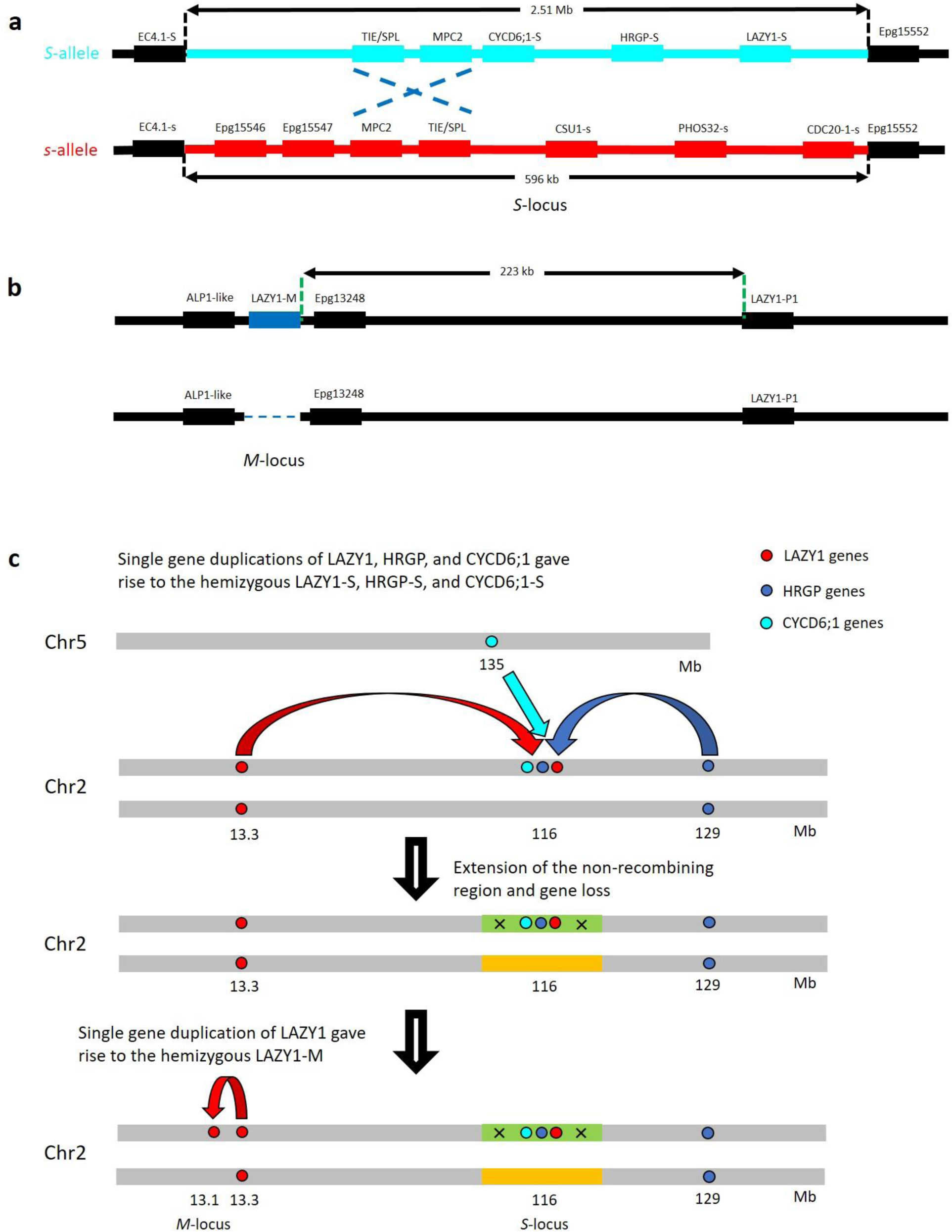
The composition and proposed model of the evolution of the *S*- and *M*-loci in *Eichhornia paniculata*. ***a,*** Genomic organization of the *S*-locus genes. Black dashed lines represent the boundaries of the tristyly alleles inferred from the read coverage analysis. Blue dashed lines represent inferred boundaries of an inversion. **b,** Genomic organization of the genes around LAZY1-M. Green dashed lines represent the boundaries of LAZY1-M and its paralog LAZY1-P1. **c,** Hypothesized evolutionary pathway for the organization of the *S*- and *M*-loci.

We hypothesized that given the high divergence between alleles, our collapsed genome assembly may have created a chimeric assembly in this region, leaving the *s*-allele sequence adjacent to the *S*-allele region in our assembled scaffold and contributing to the lack of a signal of high diversity when mapping to this reference genome. To investigate this hypothesis further, we compared our L-morph assembly (*ssmm*) to this assembly in this region using whole genome alignments. As expected, Anchorwave^29^ alignments revealed the presence of a large 2.57 Mb deletion in the *ssmm* assembly relative to the *SsMm* assembly (Fig. S4), whereas the adjacent region showed homology. Thus, we conclude that the 2.5 Mb region roughly corresponds to the *S*-allele and the 650 kb region roughly corresponds to the *s*-allele. We refined the boundaries of the two alleles using the coordinates of the closest genes in their flanking regions (Fig. 4a).

Two genes (TIE/SPL and MPC2) were shared between the *S*- and *s*-alleles. The order of the two genes are opposite in the two alleles. In the *S*-allele, both genes are on the negative strand, whereas in the *s*–allele they are on the positive strand. This finding suggests the existence of an inversion containing the two genes (Fig. 4a). The rest of the genes were allele specific: three genes (CYCD6;1-S, HRGP-S, and LAZY1-S) were specific to the *S*-allele, while five genes were specific to the *s*-allele (Fig. 4a). LAZY1 is a gene family that regulates the distribution of auxin^30^, HRGP is a superfamily of genes that forms covalently cross-linked networks in primary cell walls^31^, and CYCD6;1 is a gene that regulates the formative division of cells^32^.

### The *M*-locus contains one hemizygous gene

We identified five consecutive 50-kb windows (total length: 250 kb; genome coordinates: Chr2:13,000,000-13,250,000) that had read coverage significantly higher in the 20 *M*-morph (*ssM-*) plants than in the 20 L-morph (*ssmm*) plants (criteria: the ratio between the average read coverage of *M*-morph and L-morph plants was higher than 1.6, with the *p*-value of the *t*-test < 9.73217066 ×10^-7^, Fig. 4b). There were nine coding genes in these five windows. However, among them only one gene was missing from the *ssmm* genome assembly suggesting that it is absent from the *m*-allele (Fig. 4b). Strikingly, this absent gene, LAZY1-M, also shows strong similarity to genes in the LAZY1 family, raising the possibility that this locus acts competitively in the same pathway as LAZY1-S. Consistent with the SNP association mapping, the hemizygous *M*-locus region was located on the other side of the low-recombing region on chromosome 2 (Fig. 3e).

### Genetic degeneration at the *S*-allele

In species with supergenes, signatures of genetic degeneration, including gene loss and accumulation of TEs, are expected for the dominant haplotypes of the supergenes, because dominant haplotypes are only present in heterozygotes and thus do not exhibit recombination^33^. The *S*-allele specific gene loss (the five *s*-allele specific genes) was in accord with this prediction. In addition, we found that the density of repetitive elements in the *S*- and *M*-alleles was significantly higher than the flanking regions (*p* = 0.00001985 and *p* = 0.04731, respectively, Wilcoxon signed-rank test) (Fig. 4f and h), whereas the differences between the *s*- allele and the flanking regions were not significantly different (*p* = 0.847, Wilcoxon signed-rank test) (Fig. 4g).

### The expression pattern of hemizygous genes

Among the three *S*-allele specific genes, LAZY1-S and HRGP-S were expressed in floral organs of the *S*-morph but not non-*S*-morph plants. LAZY1-S was expressed in styles but not stamens, whereas in contrast HRGP-S was expressed in stamens but not styles (Fig. 5a). CYCD6;1-S has low expression levels in both the styles and stamens of the *S*-morph (Fig. 5a). The *M*-allele specific gene LAZY1-M was expressed in both styles and stamens (Fig. 5b).

**Figure 5.**
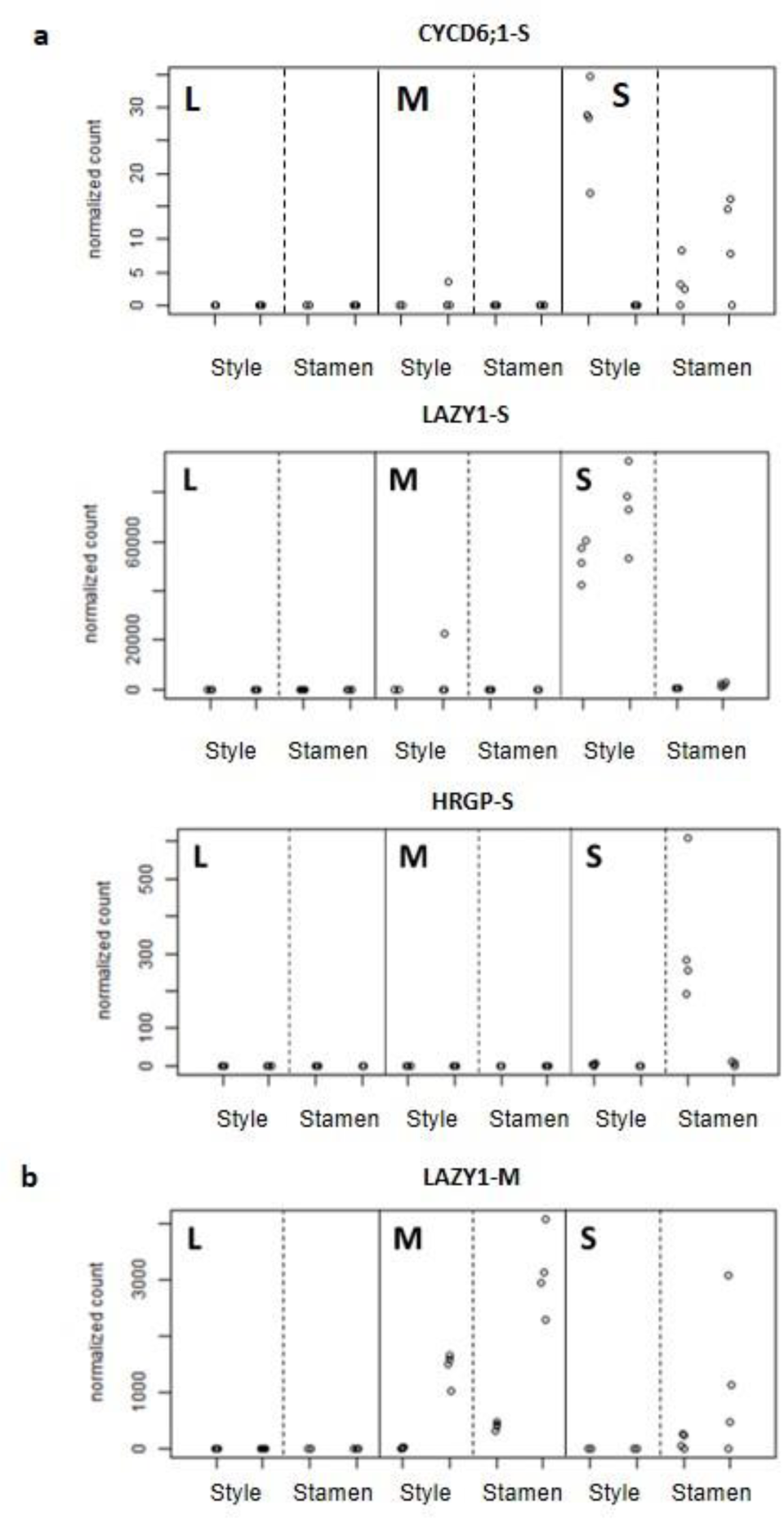
Expression patterns of the three genes that are specific to the *S*- or *M*-allele and expressed in the styles or stamens of *Eichhornia paniculata*. **a, Genes specific to the *S*-allele. b, LAZY1-M, which is specific to the *M*-allele**. Samples of different morphs are separated by solid lines. For each morph, samples of different floral organs (styles or stamens) are separated by dashed lines. For each combination of morph and floral organ, the samples at the early and late development stages are on the left and right, respectively. Each type of sample had four biological replicates.

### Genes that were differentially expressed in different morphs

For genes that were differentially expressed genome-wide between floral morphs (see Methods and Table S1 for details of comparisons), three sets of genes showed functional enrichment: (1) genes that were upregulated in *S*-morph styles (compared with non-*S*-morph styles) in early buds (enriched in three non-overlapping functions), (2) genes that were upregulated in *S*-morph styles (compared with non-*S*-morph styles) in late buds (enriched in one function), and (3) genes that were upregulated in *M*-morph styles (compared with L-morph styles) in late buds (enriched in one function) (Table S2). However, none of the five functions inferred had a direct connection to growth or flower organs, suggesting that only a small number of genes downstream to the *S*- and *M*-locus candidate genes are involved in the phenotypic differences between the floral morphs. Other differentially expressed genes may simply be incidental associations.

### The origins of the *S* and *M* loci

To further understand the origins of the *S*- and *M*-loci, we identified the closest paralogs of the *S*- and *M*-specific genes. We identified one gene that was paralogous to LAZY1-S and LAZY1-M. Hereafter, we refer to it as “LAZY1-P1”. LAZY1-P1 was located 223 kb downstream to LAZY1-M (Fig. 4b). The closest paralog of HRGP-S was found on the same chromosome but not in the same region (Fig. 4c). Two paralogs of CYCD6;1-S were annotated in both genome assemblies, and they are on chromosomes 3 and 5, respectively (hereafter CYCD6;1-Chr3 and CYCD6;1-Chr5). CYCD6;1-Chr3 shared higher sequence similarity with CYCD6;1-S (Fig. S5) and was located in a homologous region between chromosomes 2 and 3 (Fig. S2a), which was probably a result of the whole genome duplication estimated to have happened < 30 million years ago, and CYCD6;1-Chr5 is likely the ancestral copy of both CYCD6;1-S and CYCD6;1- Chr3. The distance between the three ancestral genes therefore suggests that the *S*-locus was formed via stepwise duplication rather than segmental duplication (Fig. 4c).

Based on our gene trees and assuming that the hemizygous loci represent the derived gene copies, LAZY1-S duplicated from LAZY1-P1 first, followed by subsequent gene duplication from LAZY1-P1 to form LAZY1-M. Phylogenetic dating with BEAST v2.7.6^34^ suggested that the divergence between LAZY1-S and LAZY-P1 occurred approximately 52.4 million years ago (95% confidence interval: 44.9-59.7) and that the subsequent divergence between LAZY1-M and LAZY-P1 was around 43.0 million years ago (95% confidence interval: 36.4-50.0) (Fig. 6a).

**Figure 6.**
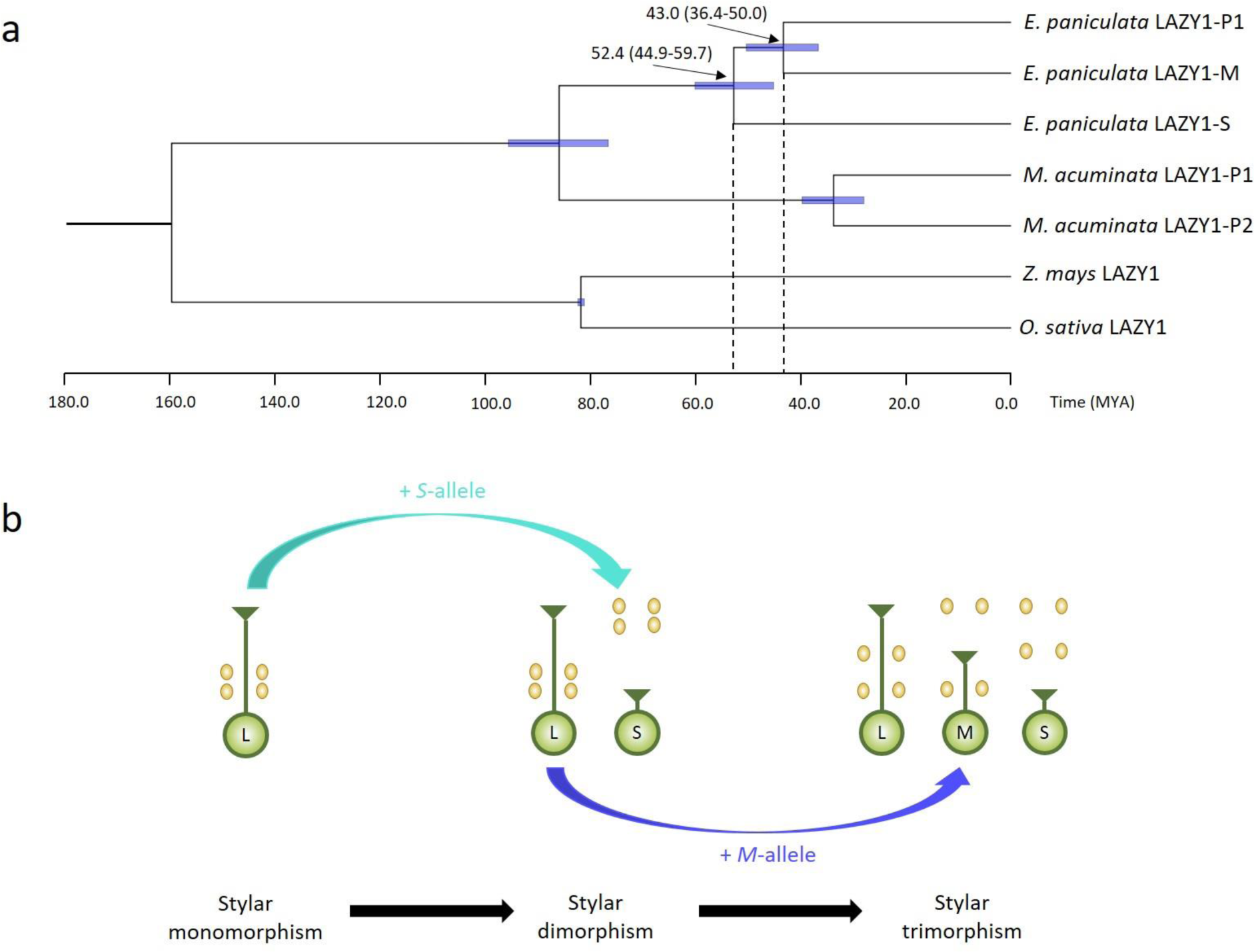
a, Phylogenetic inference of the dates of divergence between LAZY1-P1 and LAZY1-S and between LAZY1 and LAZY1-M in the ancestors of *Eichhornia paniculata*. Bayesian tree of LAZY1 genes in *Oryza sativa*, *Zea mays*, *Musa acuminata*, and *E. paniculata*. Numbers outside of parentheses represent the estimated time of divergence. Blue rectangles and numbers in parentheses are the 95% confidence intervals of the estimated time of divergence. **b, The postulated sequence of the evolution to trimorphism from monomorphism.**

## Discussion

We show that in tristylous *Eichhornia paniculata,* both the *S*- and *M*-loci are located in a low recombining region of Chromosome 2. The *S*-locus is comprised of two divergent alleles: the 2.51 Mb *S*-allele with three *S*-allele specific genes that are hemizygous in most *S*-morph plants, and the 596 kb *s*-allele with five *s*-allele specific genes. In the *S*-allele we identified two strong tristyly candidate genes: LAZY1-S and HRGP-S. LAZY1-S was expressed in styles but not stamens suggesting that it may inhibit style elongation in the *S*-morph by decreasing auxin concentration in styles. HRGP-S was expressed in stamens but not styles suggesting that it may be involved in filament elongation or possibly pollen function. Our evidence for the *S*-locus therefore strongly supports its status as a supergene, and its origin in a large region of low recombination may have been important during the initial origins of these genes via gene duplication from more physically distant regions (Figure 4).

We found that the *M*-locus contained only one hemizygous gene – LAZY1-M – and this gene was hemizygous in --*Mm* plants. Strikingly, this gene is in the same gene family as LAZY1-S suggesting the possible role of gene duplication and neofunctionalization in the origin of the epistatic interaction between the two loci. Unlike LAZY1-S, which was only expressed in styles, LAZY1-M was expressed in both styles and stamens and with a higher expression level in stamens. This suggests that while LAZY1-S may only be involved in the shortening of styles in the *S*-morph, LAZY1-M may also regulate auxin distribution and organ growth in both the styles and stamens of the *M*-morph. Given that LAZY-M is the only *M*-allele-specific hemizygous gene, the *M*-locus may not represent a supergene at this stage in its evolutionary history but simply contains LAZY1-M, a pleiotropic modifier.

LAZY1-M was also expressed in the stamens but not the styles of the *S*-morph (Fig. 5b), suggesting that LAZY1-S and LAZY1-M may compete for the same transcriptional activator and that the style expression of LAZY1-M is shut down if LAZY1-S is expressed. The expression level of LAZY1-S in *S*-morph styles is also much higher than LAZY1-M in *M*-morph styles. For example, at early development stages the average normalized count of LAZY1-M in samples of *M*-morph styles is 0.02% of that of LAZY1-S in *S*-morph styles. At late development stages, the ratio is 1.9%, indicating that the promoter region of LAZY1-S may have stronger affinity to the transcriptional activator that they compete for, potentially driving differences in style length in the *S*- and *M*-morphs through different degrees of auxin suppression.

In a theoretical treatment of the evolutionary assembly of tristyly^24^, Charlesworth predicted that the *M*-morph originated last spreading in dimorphic populations of the L- and *S*- morph and eventually giving rise to the tristylous polymorphism. Our results on the relative ages of LAZY1-S and LAZY1-M are consistent with the hypothesis that the *M*-locus evolved after the *S*-locus providing empirical support for this prediction. Our results also provide evidence suggesting a major role for ‘re-use’ through recurrent duplication of the same gene (LAZY1). Recruitment of this gene explains the origin of the *M*-locus and birth of the third floral morph in the tristylous system (Fig. 6b) resulting in the morphological and mating complexity that characterizes Darwin’s ‘complex marriage arrangement’.

## Online Methods

### Study system

Plants used in our study originated from seed collected in wild populations of *E. paniculata* in N.E. Brazil, Cuba and Mexico. The localities of plant material and details of the five populations used in our study are presented in Table S4. Information on the reproductive ecology, mating systems and population genetics of *E. paniculata* are available in Ref. 35 and references cited therein. We germinated seeds using PRO-MIX Premium Potting Mix (Premier Tech, Rivière-du-Loup, QC, Canada) and seedlings were transferred to individual 7 cm plastic pots filled with the same soil mix and placed in flooded trays in a glasshouse at the University of Toronto (Toronto, Canada) with temperatures of 25-28°C (day) and 18-21°C (night) and a 16- hour day length. Plants were then grown and flowered regularly for several years facilitating crosses and collection of leaves and floral buds for molecular analyses.

### Genome assemblies

We conducted two chromosome-level genome assemblies of *E. paniculata.* The first assembly was obtained from an individual of genotype *SsMm* (inferred from selfed progeny, Table S3) obtained from an inter-population cross between an *S*-morph plant (*SsM-*) from N.E. Brazil and an *M*-morph plant (*ssMm*) from Cuba (see Table S4 for information on the two parents). From this individual, we obtained 79 Gb (105X coverage) PacBio CLR sequences (sequenced by Arizona Genomics Institute, Tucson, AZ, USA), 55 Gb (45X coverage) Hi-C (sequenced by Phase Genomics, Seattle, WA, USA) sequences, and 117 Gb (156X coverage) 10X Chromium sequences (sequenced by The Hospital for Sick Children, Toronto, ON, Canada). The PacBio sequences were assembled with Canu^36^ and the assembly was curated with Purge Haplotigs^37^, both by the Canadian Center for Computational Genomics (C3G) (Montreal, QC, Canada). Then the genome was phased with Falcon-Phase^38^, then scaffolded into chromosome-level scaffolds with the Hi-C data, both by Phase Genomics (Seattle, WA, USA).

To identify misjoins and characterize the relationship between the physical distance and the genetic distance on each chromosome, we constructed a linkage map with genotyping-by-sequencing data obtained from a backcross pedigree from an L-morph plant and an *M*-morph plant, detailed in^25^, accessible under Accession no. SRP069136 at the Sequence Read Archive (SRA), (http://www.ncbi.nlm.nih.gov/sra, accessed 17 January 2023), and the associated BioProject alias PRJNA310331). The reads were trimmed with Scythe^39^ and Seqtk (https://github.com/lh3/seqtk). We aligned the trimmed reads to the *SsMm* genome assembly with NextGenMap V0.5.5^40^, and the BAM files were processed with Picard (http://broadinstitute.github.io/picard) using the four functions FixMateInformation, CleanSam, SortSam, and AddOrReplaceReadGroups. SAMtools^41^ was used to exclude the secondary alignments. We constructed the linkage map with the alignment files and the software program Lep-Map3^42^. Markers were split into eight large linkage groups using a logarithm of the odds (LOD) score limit of 70 and a Theta value of 0.25 for the initial split (SeparateChromosomes2). Most of the remaining single markers were put into these groups using a LOD score limit of 30 and a LOD difference of 5 (JoinSingles2All). We calculated marker order for each linkage group with OrderMarkers2 (with default settings). The first ten linkage groups contained 2,124 markers. We then used Juicebox^43^ to correct the misjoins with the linkage map. The genome was polished with the 10X Chromium sequences, using the software Pilon^44^. The repetitive elements were annotated with RepeatModeller V1.0.11^45^ and RepeatMasker V4.0.7^46^.

The second assembly of *E. paniculata* used an L-morph (*ssmm*) plant from Mexico (see Table S4) using sequencing and assembly services from Dovetail Genomics (Scotts Valley, CA, USA). 59.6 Gb (∼50X coverage) PacBio HiFi reads were generated and the reads were assembled with hifiasm^47^. The assembly was then scaffolded with Omni-C data using the HiRise software^48^. We used the linkage map previously generated to concatenate scaffolds that belong to the same linkage group. We also annotated TEs with RepeatModeller V1.0.11^45^ and RepeatMasker V4.0.7^46^. For both of our genome assemblies, the distribution of scaffold length was assessed with QUAST V5.2.0^49^, and the completeness of assemblies was assessed with BUSCO (Benchmarking Universal Single-Copy Orthologs) V5.4.4^50^ with MetaEuk^51^ and the embryophyta_odb10 (2024-01-08) dataset (Table S5).

### Genome-wide associations of single nucleotide polymorphisms (SNPs) and read coverage (RCs)

We extracted genomic DNA from 60 *E. paniculata* plants (20 of each floral morph) from a population in N.E. Brazil (hereafter “Pop 1”, see Table S4), using the QIAGEN DNeasy Plant Mini Kit (QIAGEN, Germantown, MD, USA). Library preparation and sequencing were conducted by the McGill University and Génome Québec Innovation Centre (Montréal, QC, Canada) using the 150-bp paired-end protocol on Illumina HiSeq 4000. The samples had an average sequencing depth of ∼8X.

The Illumina sequencing reads were trimmed with Scythe^39^ and Seqtk (https://github.com/lh3/seqtk). We aligned the trimmed reads to the *SsMm* genome assembly with NextGenMap V0.5.5^52^ and the BAM files were processed with Picard (http://broadinstitute.github.io/picard/) using the four functions FixMateInformation, CleanSam, SortSam, and AddOrReplaceReadGroups. We used SAMtools^41^ to exclude the secondary alignments. We used freebayes-parallel^53,54^ to call SNPs jointly for all samples. We then filtered the VCF file with VCFtools^55^ and vcflib2^56^ following a published guideline^57^. We removed sites if more than 50% of the individuals had missing data, had a minor allele count of <6, or had a quality score lower than 30. Genotypes with fewer than 4 reads were treated as “missing” and individuals with more than 50% missing data were not considered further. Then, sites with strong within-individual allelic bias at heterozygous sites (0.01 < AB < 0.25 or AB > 0.75), a discrepancy in the properly paired status for reads supporting reference or alternate alleles, a low quality relative to depth, a mean depth that was > 13, a quality score < 30, or that occurred within 5 bp of an indel, were excluded from our analysis. The resulting VCF file had 7,358,165 filtered variant sites.

We conducted a Genome-Wide Association Study (GWAS) with the variant sites using GEMMA^58^, where a relatedness matrix and a linear mixed model were used and *p*-values of variant sites were calculated with the Wald test.

To detect hemizygous regions associated with the *S*- and *M*-loci, we conducted another GWAS with read-coverage, following the pipeline in Ref. 59. We split the reference genome into 25,688 windows, each of which was 50-kb, and calculated the average depth of each window in individuals using the software mosdepth^60^. For each individual, we normalized the coverage by dividing the coverage of each window with the average coverage across all windows, then multiplied it by 2. Therefore, the expected normalized coverage is 1 for hemizygous regions and 2 for the rest of the genome. We conducted two *t*-tests for normalized coverage for each window: one for the comparison between *S*-morph versus non-*S*-morph plants, and the other for the comparison between the *M*-morph versus L-morph plants. We identified tristyly allele-specific windows with the following criteria:

1. *p* value of the *t*-test was smaller than 9.73217066 ×10^-7^, i.e. Bonferroni correction with α = 0.05.
2. We set thresholds on the ratio between the average read coverage of plants of different morphs:

For *S*-allele specific windows: The read coverage of a *S*-allele is at least 4 times of a *s*- allele, i.e. the ratio between the average read coverage of non-*S*-morph plants and *S*-morph plants was lower than 0.4: 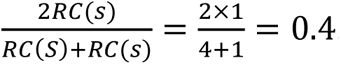

For *s*-allele specific windows: The read coverage of a *s*-allele is at least 4 times of a *S*-allele, i.e. the ratio between the average read coverage of non-*S*-morph plants and *S*-morph plants was higher than 1.6: 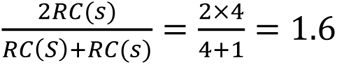

For *M*-allele specific windows: The read coverage of an *M*-allele is at least 4 times of an *m*-allele, i.e. the ratio between the average read coverage of non-M-morph plants and *M*-morph plants was lower than 0.25, assuming all *M*-morph plants were homozygous: 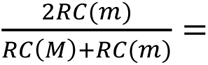 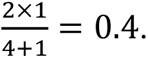

For *m*-allele specific windows: The read coverage of an *s*-allele is at least 4 times of a *M*-allele, i.e. the ratio between the average read coverage of non-M-morph plants and *M*-morph plants was higher than 1.6: 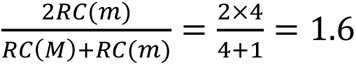

Only regions with at least two consecutive windows that were allele-specific were considered. As a result, a 2.5 Mb *S*-allele, a 650 kb *s*-allele, and a 250 kb *M*-allele were identified. We split each allele into 10 equal-sized windows (window sizes: *S*-allele: 250 kb; *s*-allele: 65 kb; *M*-allele: 25 kb), calculated the repeat content of each window, then compared it with the 10 windows of the same size flanking each allele (5 windows upstream and 5 windows downstream of the locus) with Wilcoxon rank-sum tests.

### Differential gene expression analysis and genome annotation with transcriptome data

To infer genes controlling different components of the tristyly syndrome, we conducted a floral organ-specific transcriptomic analysis. We dissected floral buds of the three morphs at two developmental stages: “early”, with buds between 3-5 mm in length, and “late”, with buds between 5-8 mm. We used sterilized forceps under a stereo dissecting microscope to open the floral buds and placed the styles and stamens into 2 mL centrifuge tubes which were then frozen by putting them into liquid nitrogen. We extracted RNA from the frozen tissues with Spectrum™ Plant Total RNA-Kit (Sigma-Aldrich, St. Louis, MO, USA) following the manufacturer’s instructions. For each type of sample, we collected four biological replicates, each containing styles or stamens from 5-27 buds sampled from 5-17 individuals from each of two populations (Pop. 1, 2) originating from N.E. Brazil (Table S4, S6).

We prepared sequencing libraries and sequenced on the Illumina HiSeq4000 platform (100-bp paired-end reads) by the McGill University and Génome Québec Innovation Centre (Montréal, QC, Canada). Then we aligned the Illumina sequences to the *SsMm* genome assembly with HISAT2^61^ and assembled the transcripts with StringTie^62^. We conducted functional annotations of the floral organ transcriptome with Mercator4^63^. We used StringTie^62^ to estimate the abundance of each transcript in each sample. Then we used the R package DESeq2^64^ to conduct differential gene expression analysis and plot the normalized count in each sample for the genes at the *S*- and *M*-loci. We conducted eight differential gene expression analyses for each type of morph difference (*S* vs. non-S and M vs. L), each tissue (style vs. stamen), and each development stage (early bud vs. late bud). Only genes with FPKM (Fragments Per Kilobase of transcript per Million mapped reads) > 1 and TPM (Transcript Per Million) > 1 in at least one sample were included in the differential gene expression analyses. In each analysis, we identified the genes that are upregulated in each morph with false discovery rate < 0.05 and LFC (log fold change) > 2 or < −2. We then did a Mercator BIN enrichment analysis^61^ with the list of upregulated genes for each comparison.

We also annotated the repeat-masked versions of the two genome assemblies with the floral organ transcriptomes and also leaf and floral bud transcriptomes from Ref. 65, available under accession number SRP049636 on the Sequence Read Archive (SRA) (http://www.ncbi.nlm.nih.gov/sra, accessed 20 January 2023) and the associated BioProject alias is PRJNA266681 (http://www.ncbi.nlm.nih.gov/bioproject/, accessed 20 January 2023), and also Ref. 66, available under accession number SRP060405 at the Sequence Read Archive and associated BioProject alias (PRJNA288861) using MAKER v3.01.03^67^ and InterProScan 5^68^. This resulted in a total of 35,224 annotated genes for the *SsMm* genome and 29,592 genes for the *ssmm* genome.

### Origin of the *S* and *M* loci

According to the annotation of the *SsMm* and *ssmm* genome assemblies, there is only one LAZY1 homolog (LAZY1-P1) that exists in both genome assemblies. We then downloaded the coding sequences of LAZY1 in *Oryza sativa* and *Zea mays* from NCBI (https://www.ncbi.nlm.nih.gov/, accession numbers: NM_001138862.1 and DQ855268.1), and two paralogs of LAZY1 in *Musa acuminata* ‘DH-Pahang’ from the website Banana Genome Hub (https://banana-genome-hub.southgreen.fr/). We conducted a multiple sequence alignment of these sequences and the coding sequences of three LAZY1 genes in *E. paniculata*, using MAFFT v7^69^ with L-IN*S*-i method.

We then estimated the time of divergence between LAZY1-S and LAZY1-P1 and between LAZY1-M and LAZY1-P1 with BEAST v2.7.6^34^. First, we did model selection of jModelTest^70^ using Akaike information criterion (AIC) as the criteria. The GTR + I + Gamma model with nCat = 4 was selected. Then we set the time to the most recent ancestors of *Oryza sativa* and *Zea mays* (81.43 ±0.3 MYA)^71^, Zingiberales and Commelinales (79.8 ±9.87 MYA)^72^, and Poales and Commelinales (106.7 ±8.3 MYA)^72^ as calibration points. Ten independent Markov Chain Monte Carlo runs, each with 1 x 10^8^ generations (first 10% was burn-in) were conducted. We combined the output of the eight runs using LogCombiner v2.7.6^34^ with a sample frequency of 5,000. Then we used TreeAnnotator v2.7.6^34^ to generate the maximum clade credibility tree, which was visualized in FigTree v1.4.4 (http://tree.bio.ed.ac.uk/software/figtree/) (Fig. 6).

There are two CYCD6;1 paralogs that exists in both genome assemblies, and they are on chromosomes 3 and 5, respectively (hereafter CYCD6;1-Chr3 and CYCD6;1-Chr5). We then downloaded the coding sequence of CYCD6;1 in *O. sativa* from NCBI (https://www.ncbi.nlm.nih.gov/, accession numbers: XM_015789871.2). We conducted a multiple sequence alignment of the coding sequences of the three CYCD6;1 genes using MAFFT v7^69^ with the L-IN*S*-i method. Then we constructed a Neighbor Joining tree with all the conserved sites, Jukes-Cantor substitution model, and bootstrap resampling with 100 samples. (Fig. S).

HRGP-S was not annotated by MAKER or InterProScan. We aligned the transcript of HRGP-S against the *SsMm* and *ssmm* genome assemblies with the software program blastn^73^, to identify its closest paralog.

### Synteny analyses, whole genome alignments and structural variants

We used COGE’s SynMap2 algorithm^74^ with the MAKER and InterProScan annotations to examine the synteny between the two genome assemblies and identify structural variation. To obtain finer-scale whole genome alignment of syntenic regions to localize putative insertion/deletion differences between the *S*- and *s*-alleles and between the *M*- and *m*-alleles, we used Anchorwave^29^ to align the two genome assemblies, with the “minimap2” option and the “proali” function.

### Population genomic statistics

We calculated the nucleotide diversity (*π*) of 1-Mb sliding windows for the 60 individuals (20 of each morph) used in the GWAS analyses. We used SAMtools^41^ to remove the reads with mapping quality < 10 in the filtered BAM files generated in the section “Genome-wide associations of single nucleotide polymorphisms (SNPs) and read coverage (RCs)”. Then we used ANGSD^75^ to calculate the nucleotide diversity (*π*) of the sliding windows.

## Acknowledgements

This work was supported by Discovery Grants from the Natural Sciences and Engineering Research Council of Canada (NSERC) to S.C.H.B. and S.I.W. and student fellowships from the University of Toronto to H.X. The Canu assembly and Purge Haplotigs pipeline of the *SsMm* genome was conducted by the Canadian Center for Computational Genomics (C3G), which is a Genomics Technology Platform (GTP) supported by the Canadian Government through Genome Canada. We thank William Cole and Tom Gludovacz for glasshouse maintenance, Joanna Tran, SungA Choe, Ryan Ruan, and Jianuo Feng for help with growing plants, Claire Ellis and Po-Yuan Huang for help with tissue collection, Baharul Choudhury, Iuliia Pelikh, and Yihe Zheng for help with DNA extraction, and Joanna Rifkin, Christian Kappel, Mathias Scharmann, Julia Kreiner, Jingshan Yang, Meng Yuan, Liangjiao Xue, ZoëHumphries, Tyler Kent, and Ting Liu for advice on data analysis.

## Author contributions

H.X., S.I.W., and S.C.H.B. conceived and designed the study and wrote the manuscript. S.C.H.B. obtained the seeds used for the study. H.X. grew the plants, collected the tissues, and extracted DNA and RNA. Y.G. performed the genome annotation. S.I.W. performed the Anchorwave alignment. H.X. performed all remaining analyses. H.X., S.I.W., and S.C.H.B. wrote the manuscript. All authors read and approved the final manuscript.

## Data Availability Statement

The PacBio CLR sequences used to assemble the *Eichhornia paniculata SsMm* genome are available under Accession no. SRR21038607 at the Sequence Read Archive (SRA) (http://www.ncbi.nlm.nih.gov/sra, accessed 09 February 2024), and the associated BioProject alias is PRJNA863567. The Hi-C data used to assemble the *E. paniculata SsMm* genome are available under Accession no. SRR21038606 at the Sequence Read Archive (SRA) (http://www.ncbi.nlm.nih.gov/sra, accessed 09 February 2024), and the associated BioProject alias is PRJNA863567. The 10X Chromium data used to polish the *E. paniculata SsMm* genome are available under Accession no. SRR21038605 at the Sequence Read Archive (SRA) (http://www. ncbi.nlm.nih.gov/sra, accessed 09 February 2024), and the associated BioProject alias is PRJNA863567. The PacBio CCS sequences used to assemble the *E. paniculata ssmm* genome are available under Accession no. SRRXXXXXXXX at the Sequence Read Archive (SRA) (http://www.ncbi.nlm.nih.gov/sra, accessed XX XX XXXX), and the associated BioProject alias is PRJNAXXXXXX. The Omni-C data used to assemble the *E. paniculata ssmm* genome are available under Accession no. SRRXXXXXX at the Sequence Read Archive (SRA) (http://www.ncbi.nlm.nih.gov/sra, accessed XX XX XXXX), and the associated BioProject alias is PRJNAXXXXXX. The whole genome Illumina sequences of the 60 *E. paniculata* individuals in this study are available under Accession no. SRRXXXXXXXX - SRRXXXXXXXX at the Sequence Read Archive (SRA) (http://www.ncbi.nlm.nih.gov/sra, accessed XX XX XXXX), and the associated BioProject alias is PRJNA862356. The RNA-Seq data of styles and stamens are available under Accession no. SRRXXXXXXXX - SRRXXXXXXXX at the Sequence Read Archive (SRA) (http://www.ncbi.nlm.nih.gov/sra, accessed XX XX XXXX), and the associated BioProject alias is PRJNAXXXXXX. The *E. paniculata SsMm* genome and its MAKER and InterProScan annotation are available under Genome ID XXXXX on CoGe (https://genomevolution.org/coge/, accessed XX XX XXXX). The *E. paniculata ssmm* genome and its MAKER and InterProScan annotation are available under Genome ID XXXXX on CoGe (https://genomevolution.org/coge/, accessed XX XX XXXX).

## References

1. Darwin, C. R. The different forms of flowers on plants of the same species. (John Murray, 1877).

2. Barrett, S. C. H. Heterostylous genetic polymorphisms: model systems for evolutionary analysis. In Evolution and function of heterostyly (ed. Barrett, S. C. H.) 1–29 (Springer, 1992).

3. Barrett, S. C. H. The evolution of plant sexual diversity. Nat. Rev. Genet. 3, 274–284 (2002).

4. Lloyd, D. G. & Webb, C. The selection of heterostyly. In Evolution and function of heterostyly (ed. Barrett, S. C. H.) 179–207 (Springer, 1992).

5. Lloyd, D. G. & Webb, C. J. The evolution of heterostyly. In Evolution and function of heterostyly (ed. Barrett, S. C. H.) 151–178 (Springer, 1992).

6. Barrett, S. C. H., Jesson, L. K. & Baker, A. M. The evolution and function of stylar polymorphisms in flowering plants. Ann. Bot. 85, 253–265 (2000).

7. Lewis, D. & Jones, D. The genetics of heterostyly. In Evolution and function of heterostyly (Ed. Barrett, S. C. H.) 129–150 (Springer, 1992).

8. Mather, K. The genetical architecture of heterostyly in *Primula sinensis*. Evolution 4, 340–352 (1950).

9. Schwander, T., Libbrecht, R. & Keller, L. Supergenes and complex phenotypes. Current Biology 24, 288–294 (2014).

10. Thompson, M. J. & Jiggins, C. Supergenes and their role in evolution. Heredity 113, 1–8 (2014).

11. Maney, D. L. & Küpper, C. Supergenes on steroids. Philos. Trans. R. Soc. B 377, 20200507 (2022).

12. Kappel, C., Huu, C. N. & Lenhard, M. A short story gets longer: recent insights into the molecular basis of heterostyly. J. Exp. Bot. 68, 5719–5730 (2017).

13. Barrett, S. C. H. ‘A most complex marriage arrangement’: recent advances on heterostyly and unresolved questions. New Phytol. 224, 1051–1067 (2019).

14. Huu, C. N. et al. Presence versus absence of CYP734A50 underlies the style-length dimorphism in primroses. eLife 5, e17956 (2016).

15. Li, J. et al. Genetic architecture and evolution of the S locus supergene in *Primula vulgaris*. Nat. Plants 2, 1–7 (2016).

16. Shore, J. S. et al. The long and short of the *S*-locus in *Turnera* (Passifloraceae). New Phytol. 224, 1316–1329 (2019).

17. Gutiérrez-Valencia, J. et al. Genomic analyses of the *Linum* distyly supergene reveal convergent evolution at the molecular level. Curr. Biol. 32, 4360–4371 (2022).

18. Zhao, Z. et al. Genomic evidence supports the genetic convergence of a supergene controlling the distylous floral syndrome. New Phytol. 237, 601–614 (2023).

19. Yang, J. et al. Haplotype-resolved genome assembly provides insights into the evolution of *S*-locus supergene in distylous *Nymphoides indica*. New Phytol. 240, 2058–2071 (2023).

20. Fawcett, J. A. et al. Genome sequencing reveals the genetic architecture of heterostyly and domestication history of common buckwheat. Nature Plants 9, 1236–1251 (2023).

21. Potente, G. et al. Comparative genomics elucidates the origin of a supergene controlling floral heteromorphism. Mol. Biol. Evol. 39, msac035 (2022).

22. Barrett, S. C. H. The evolutionary biology of tristyly. In Oxford Surveys in Evolutionary Biology (Eds. D. Futuyma & J. Antonovics) vol. 9, 283–326 (Oxford University Press, 1993).

23. Thompson, J. D., Pailler, T., Strasberg, D. & Manicacci, D. Tristyly in the endangered Mascarene Island endemic *Hugonia serrata* (Linaceae). Am. J. Bot. 83, 1160–1167 (1996).

24. Charlesworth, D. The evolution and breakdown of tristyly. Evolution 33, 489–498. (1979)

25. Arunkumar, R., Wang, W., Wright, S. I. & Barrett, S. C. H. The genetic architecture of tristyly and its breakdown to self-fertilization. Mol. Ecol. 26, 752–765 (2017).

26. Ossowski, S. et al. The rate and molecular spectrum of spontaneous mutations in *Arabidopsis thaliana*. Science 327, 92–94 (2010).

27. Eckenwalder, J. E. & Barrett, S. C. H. Phylogenetic systematics of Pontederiaceae. Syst. Bot. 11, 373–391 (1986).

28. Ness, R. W., Siol, M. & Barrett, S. C. H. *De novo* sequence assembly and characterization of the floral transcriptome in cross-and self-fertilizing plants. BMC Genomics 12, 298 (2011).

29. Song, B. et al. AnchorWave: Sensitive alignment of genomes with high sequence diversity, extensive structural polymorphism, and whole-genome duplication. Proc. Natl. Acad. Sci. 119, e2113075119 (2022).

30. Jiao, Z., Du, H., Chen, S., Huang, W. & Ge, L. LAZY gene family in plant gravitropism. Front. Plant Sci. 11, 606241 (2021).

31. Chen, Y. et al. Identification of the abundant hydroxyproline-rich glycoproteins in the root walls of wild-type *Arabidopsis*, an ext3 mutant line, and its phenotypic revertant. Plants 4, 85– 111 (2015).

32. Sozzani, R. et al. Spatiotemporal regulation of cell-cycle genes by SHORTROOT links patterning and growth. Nature, 466, 128–132 (2021).

33. Gutiérrez-Valencia, J. et al. The genomic architecture and evolutionary fates of supergenes. Genome Biology and Evolution 13, evab057 (2021).

34. Bouckaert, R. et al. BEAST 2.5: An advanced software platform for Bayesian evolutionary analysis. PLoS Comput. Biol. 15, e1006650 (2019).

35. Ness, R. W., Wright, S. I. & Barrett, S. C. H. Mating-system variation, demographic history and patterns of nucleotide diversity in the tristylous plant *Eichhornia paniculata*. Genetics 184, 381–392 (2010).

36. Koren, S. et al. Canu: scalable and accurate long-read assembly via adaptive k-mer weighting and repeat separation. Genome Res. 27, 722–736 (2017).

37. Roach, M. J., Schmidt, S. A., & Borneman, A. R. Purge Haplotigs: allelic contig reassignment for third-gen diploid genome assemblies. BMC bioinformatics 19, 460 (2018).

38. Kronenberg, Z. N., et al. Extended haplotype-phasing of long-read de novo genome assemblies using Hi-C. Nature Communications 12, 1935 (2021).

39. Thress, K., Henzel, W., Shillinglaw, W. & Kornbluth, S. Scythe: a novel reaper-binding apoptotic regulator. EMBO J. 17, 6135–6143 (1998).

40. Sedlazeck, F. J., Rescheneder, P. & Von Haeseler, A. NextGenMap: fast and accurate read mapping in highly polymorphic genomes. Bioinformatics 29, 2790–2791 (2013).

41. Danecek, P. et al. Twelve years of SAMtools and BCFtools. GigaScience 10, (2021).

42. Rastas, P. Lep-MAP3: robust linkage mapping even for low-coverage whole genome sequencing data. Bioinformatics 33, 3726–3732 (2017).

43. Durand, N. C. et al. Juicebox provides a visualization system for Hi-C contact maps with unlimited zoom. Cell Syst. 3, 99–101 (2016).

44. Walker, B. J. et al. Pilon: an integrated tool for comprehensive microbial variant detection and genome assembly improvement. PLoS One 9, e112963 (2014).

45. Smit, A. F. A. & Hubley, R. RepeatModeler Open-1.0. 2008–2015. (2008).

46. Smit, A. F. A., Hubley, R. & Green, P. RepeatMasker Open-4.0. (2013).

47. Cheng, H., Concepcion, G. T., Feng, X., Zhang, H. & Li, H. Haplotype-resolved de novo assembly using phased assembly graphs with hifiasm. Nat. Methods 18, 170–175 (2021).

48. Putnam, N. H. et al. Chromosome-scale shotgun assembly using an *in vitro* method for long-range linkage. Genome Res. 26, 342–350 (2016).

49. Gurevich, A., Saveliev, V., Vyahhi, N. & Tesler, G. QUAST: quality assessment tool for genome assemblies. Bioinformatics 29, 1072–1075 (2013).

50. Manni, M. et al. BUSCO update: novel and streamlined workflows along with broader and deeper phylogenetic coverage for scoring of eukaryotic, prokaryotic, and viral genomes. Mol. Biol. Evol. 38, 4647–4654 (2021).

51. Levy Karin, E., Mirdita, M., & Söding, J. MetaEuk—sensitive, high-throughput gene discovery, and annotation for large-scale eukaryotic metagenomics. Microbiome 8, 48 (2020).

52. Sedlazeck, F. J., Rescheneder, P. & Von Haeseler, A. NextGenMap: fast and accurate read mapping in highly polymorphic genomes. Bioinformatics 29, 2790–2791 (2013).

53. Garrison, E. & Marth, G. Haplotype-based variant detection from short-read sequencing. ArXiv Prepr., ArXiv12073907 (2012).

54. Tange, O. Gnu parallel: the command-line power tool. Usenix Mag. 36, 42 (2011).

55. Danecek, P. et al. The variant call format and VCFtools. Bioinformatics 27, 2156–2158 (2011).

56. Garrison, E., Kronenberg, Z. N., Dawson, E. T., Pedersen, B. S. & Prins, P. A spectrum of free software tools for processing the VCF variant call format: vcflib, bio-vcf, cyvcf2, hts-nim and slivar. PLoS Comput. Biol. 18, e1009123 (2022).

57. Puritz, J. B., Hollenbeck, C. M. & Gold, J. R. dDocent: a RADseq, variant-calling pipeline designed for population genomics of non-model organisms. PeerJ 2, e431 (2014).

58. Zhou, X. & Stephens, M. Genome-wide efficient mixed-model analysis for association studies. Nat. Genet. 44, 821–824 (2012).

59. Xue, L. et al. Evidences for a role of two Y-specific genes in sex determination in *Populus deltoides*. Nature Communications 11, 5893 (2020).

60. Pedersen, B. S. & Quinlan, A. R. Mosdepth: quick coverage calculation for genomes and exomes. Bioinformatics 34, 867–868 (2018).

61. Kim, D., Langmead, B. & Salzberg, S. L. HISAT: a fast spliced aligner with low memory requirements. Nat. Methods 12, 357–360 (2015).

62. Pertea, M. et al. StringTie enables improved reconstruction of a transcriptome from RNA-seq reads. Nat. Biotechnol. 33, 290–295 (2015).

63. Lohse, M., et al. Mercator: a fast and simple web server for genome scale functional annotation of plant sequence data. Plant, Cell & Environment 37, 1250–1258 (2014).

64. Love, M. I., Huber, W. & Anders, S. Moderated estimation of fold change and dispersion for RNA-seq data with DESeq2. Genome Biol. 15, 1–21 (2014).

65. Arunkumar, R., Ness, R. W., Wright, S. I. & Barrett, S. C. H. The evolution of selfing is accompanied by reduced efficacy of selection and purging of deleterious mutations. Genetics 199, 817–829 (2015).

66. Arunkumar, R., Maddison, T. I., Barrett, S. C. H. & Wright, S. I. Recent mating-system evolution in *Eichhornia* is accompanied by cis-regulatory divergence. New Phytol. 211, 697–707 (2016).

67. Cantarel, B. L. et al. MAKER: an easy-to-use annotation pipeline designed for emerging model organism genomes. Genome Res. 18, 188–196 (2008).

68. Jones, P. et al. InterProScan 5: genome-scale protein function classification. Bioinformatics 30, 1236–1240 (2014).

69. Katoh, K. & Standley, D. M. MAFFT Multiple Sequence Alignment Software Version 7: Improvements in Performance and Usability. Mol. Biol. Evol. 30, 772–780 (2013).

70. Posada, D. jModelTest: Phylogenetic Model Averaging. Mol. Biol. Evol. 25, 1253–1256 (2008).

71. Huang, W. et al. A well-supported nuclear phylogeny of Poaceae and implications for the evolution of C4 photosynthesis. Mol. Plant 15, 755–777 (2022).

72. Magallón, S., Gómez-Acevedo, S., Sánchez-Reyes, L. L. & Hernández-Hernández, T. A metacalibrated time-tree documents the early rise of flowering plant phylogenetic diversity. New Phytol. 207, 437–453 (2015).

73. Camacho, C. et al. BLAST+: architecture and applications. BMC Bioinformatics 10, 421 (2009).

74. Haug-Baltzell, A., Stephens, S. A., Davey, S., Scheidegger, C. E. & Lyons, E. SynMap2 and SynMap3D: web-based whole-genome synteny browsers. Bioinformatics 33, 2197–2198 (2017).

75. Korneliussen, T. S., Albrechtsen, A., & Nielsen, R. ANGSD: analysis of next generation sequencing data. BMC bioinformatics 15, 356 (2014).

