## Supplementary Materials for "Genomic evidence for supergene control of Darwin’s “complex marriage arrangement” – the tristylous floral polymorphism"

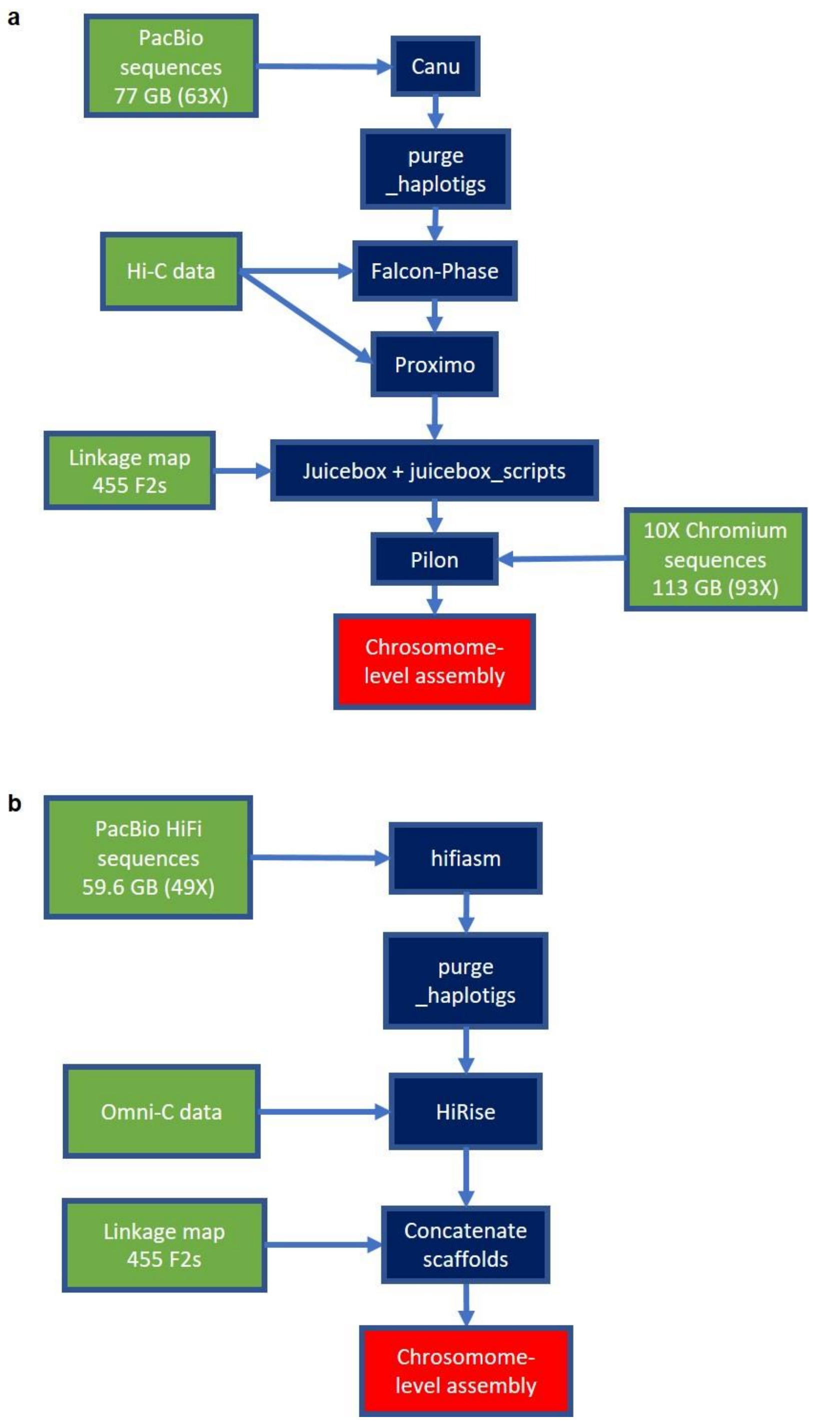


**Figure S1 | Schematics of assembly workflow.** a. *SsMm* assembly. b. *ssmm* assembly for *Eichhornia paniculata*

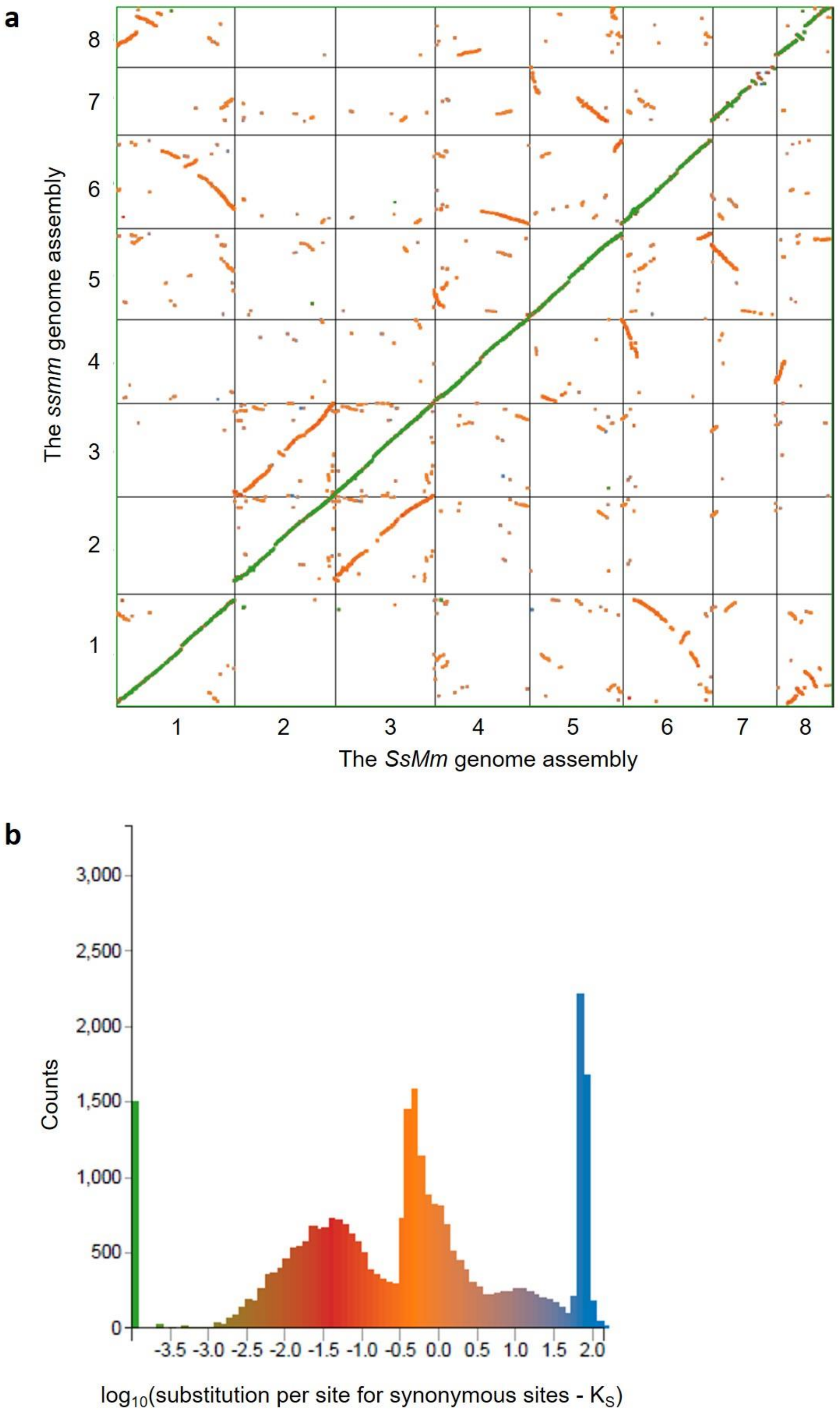
a
8
7
6
5
4
3
2
1
The ssmm genome assembly
1
2
3
4
5
6
7
8
The SsMm genome assembly
b
3,000
2,500
2,000
1,500
1,000
500
0
Counts
-3.5 -3.0 -2.5 -2.0 -1.5 -1.0 -0.5 0.0 0.5 1.0 1.5 2.0
log10(substitution per site for synonymous sites - KS)

**Figure S2 | a.** Syntenic comparison between the *SsMm* (x-axis) and *ssmm* (y-axis) assemblies of *Eichhornia paniculata*. Colours represent synonymous substitution rates of homologous genes shown in b. **b.** Distribution of synonymous substitutions between homologous genes.

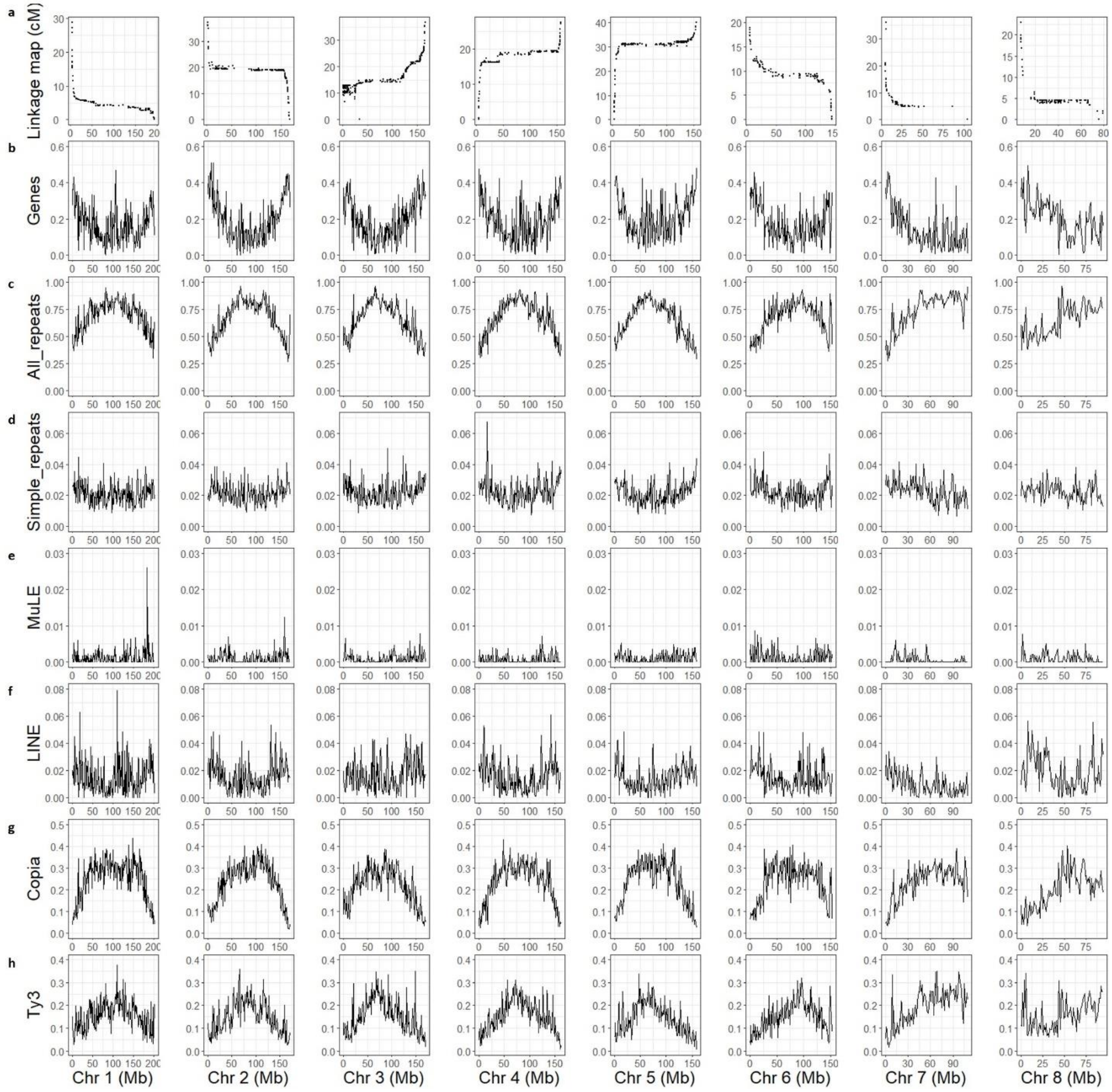

**Figure S3 | The recombination landscape and genome content across the chromosomes of *Eichhornia paniculata*. a**. Marey maps plotting the position on the linkage map (cM) against the position on the physical map (Mb). **b-h,** Densities of genes (**b**), all repetitive elements (**c**), simple repeats (**d**), Mutator-Like Element (MuLE) (**e**), Long Interspersed Nuclear Element (LINE) (**f**), Copia elements (**g**), and Ty3 elements (**h**), all measured as proportions of 1-Mb windows.

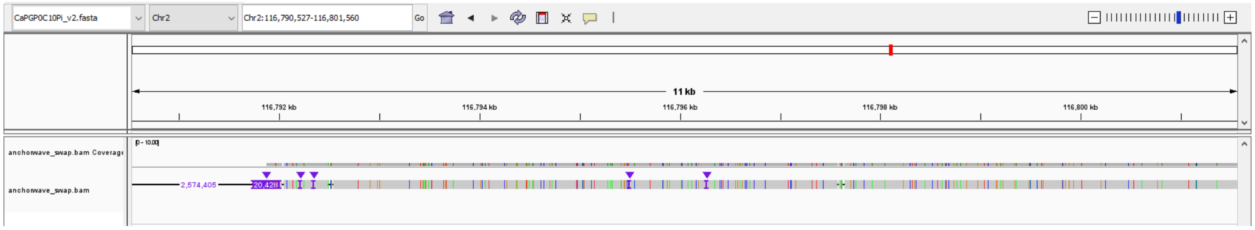


**Figure S4 | Anchorwave alignment supports divergence between the *S* and *s* alleles in *Eichhornia paniculata*.**

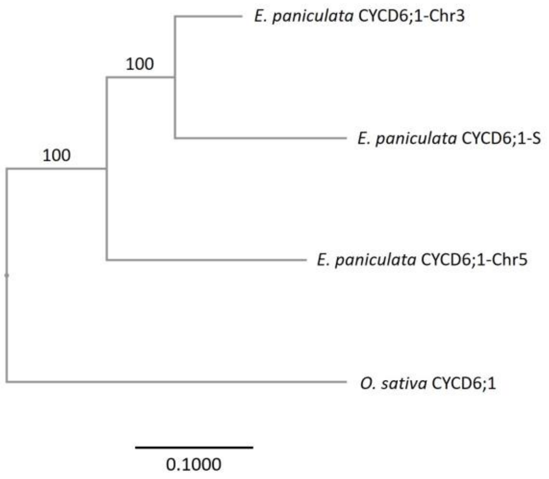

**Figure S5 | Neighbor joining tree of CYCD6;1 genes in *Oryza sativa* and *Eichhornia paniculata*.** Branch length represents substitutions per site. Numbers on branches are bootstrap values.

**Table S1 | Number of differentially expressed genes in each comparison between the morphs of *Eichhornia paniculata***

| Comparison | | | LFC > 2 | LFC < -2 |
|---|---|---|---|---|
| Morph | Tissue | Development stage | | |
| S vs. non-S | Style | Early | 450 | 586 |
| S vs. non-S | Style | Late | 854 | 609 |
| S vs. non-S | Stamen | Early | 413 | 406 |
| S vs. non-S | Stamen | Late | 485 | 418 |
| M vs. L | Style | Early | 704 | 775 |
| M vs. L | Style | Late | 353 | 292 |
| M vs. L | Stamen | Early | 781 | 374 |
| M vs. L | Stamen | Late | 219 | 138 |

Genes that were significantly (adjusted $p < 0.05$) upregulated (LFC > 2) or downregulated (LFC < -2) in S-morph plants (compared to non-S-morph plants) or M-morph plants (compared to L-morph plants) in styles or stamens at early (bud size = 3-5 mm) or late (bud size = 5-8 mm) development stages.

**Table S2 | Comparisons with significant enrichments in Mercator4 BIN enrichment analysis**

Upregulated in S-morph styles (compared with non-S-morph styles), early developmental stage

| MapMan4 category number | Context of Protein Function | #Genes of Interest IN MapMan4 category | #Genes of Interest NOT IN MapMan4 category | #Background Genes IN MapMan4 category | #Background Genes NOT IN MapMan4 category | Enrichment Factor | p-value | FDR-adjusted p-value |
|---|---|---|---|---|---|---|---|---|
| 11.7.3.3 | Phytohormone action.jasmonic acid.conjugation and degradation.jasmonoyl-amino acid carboxylase *(CYP94C) | 4 | 446 | 5 | 38178 | 67.88 | 9.430282226897081e-8 | 0.00016958005014517676 |
| 25.2.4.2 | Nutrient uptake.sulfur assimilation.sulfate homeostasis.regulatory protein *(LSU) | 3 | 447 | 5 | 38178 | 50.91 | 0.000015977266307294948 | 0.018968594915591348 |
| 11.7.3 | Phytohormone action.jasmonic acid.conjugation and degradation | 7 | 443 | 16 | 38167 | 37.12 | 3.1465123530129466e-10 | 7.544287785074041e-7 |
| 11.7 | Phytohormone action.jasmonic acid | 7 | 443 | 52 | 38131 | 11.42 | 0.0000025518947712634l6 | 0.0036711558179395505 |
| 50.1.13 | Enzyme classification.EC_1 oxidoreductases.EC_1.14 oxidoreductase acting on paired donor with incorporation or reduction of molecular oxygen | 13 | 437 | 265 | 37918 | 4.16 | 0.000018459636369962382 | 0.018968594915591348 |
| 35 | not assigned | 326 | 124 | 19558 | 18625 | 1.41 | 2.0257147022604762e-20 | 7.285482926679802e-17 |
| 35.2 | not assigned.not annotated | 326 | 124 | 19558 | 18625 | 1.41 | 2.0257147022604762e-20 | 7.285482926679802e-17 |

Upregulated in S-morph styles (compared with non-S-morph styles), late developmental stage

| MapMan4 category number | Context of Protein Function | #Genes of Interest IN MapMan4 category | #Genes of Interest NOT IN MapMan4 category | #Background Genes IN MapMan4 category | #Background Genes NOT IN MapMan4 category | Enrichment Factor | p-value | FDR-adjusted p-value |
|---|---|---|---|---|---|---|---|---|
| 14.4.4 | DNA damage response.nonhomologous end-joining (NHEJ) repair.protein ADP-ribosyltransferase *(PARP3) | 3 | 773 | 3 | 38180 | 49.2 | 0.000008362350464846565 | 0.020050128964547113 |
| 35 | not assigned | 610 | 166 | 19558 | 18625 | 1.53 | 3.9433201059744084e-57 | 1.418215076113696e-53 |
| 35.2 | not assigned.not annotated | 610 | 166 | 19558 | 18625 | 1.53 | 3.9433201059744084e-57 | 1.418215076113696e-53 |

Upregulated in M-morph styles (compared with L-morph styles), late developmental stage

| MapMan4 category number | Context of Protein Function | #Genes of Interest IN MapMan4 category | #Genes of Interest NOT IN MapMan4 category | #Background Genes IN MapMan4 category | #Background Genes NOT IN MapMan4 category | Enrichment Factor | p-value | FDR-adjusted p-value |
|---|---|---|---|---|---|---|---|---|
| 9.2.2.2.1.1 | Secondary metabolism.phenolics.flavonoid biosynthesis.chalcones.chalcone synthase activity.chalcone synthase *(CHS) | 6 | 848 | 11 | 38172 | 24.39 | 5.163993282548907e-8 | 0.00012381534560458094 |
| 9.2.2.2 | Secondary metabolism.phenolics.flavonoid biosynthesis.chalcones | 6 | 848 | 18 | 38165 | 14.9 | 0.000001814511652293108 | 0.002610356462988865 |
| 9.2.2.2.1 | Secondary metabolism.phenolics.flavonoid biosynthesis.chalcones.chalcone synthase activity | 6 | 848 | 18 | 38165 | 14.9 | 0.000001814511652293108 | 0.002610356462988865 |
| 35 | not assigned | 605 | 249 | 19558 | 18625 | 1.38 | 3.194365884954564e-32 | 1.1488536905239088e-28 |
| 35.2 | not assigned.not annotated | 605 | 249 | 19558 | 18625 | 1.38 | 3.194365884954564e-32 | 1.1488536905239088e-28 |

***Comparisons for which only "35, not assigned" and "35.2, not assigned.not annotated" were significantly enriched are not shown.**

**Table S3 | Floral morphs of individuals in the selfed progeny of the S-morph plant (*SsMm*) used for genome assembly**

| Floral morph | Number of individuals | Percentage |
|---|---|---|

| | | |
|---|---|---|
| L-morph | 117 | 21.6% |
| M-morph | 9 | 1.7% |
| S-morph | 415 | 76.7% |
| Total | 541 | |

**Table S4 | Information on the locations of *Eichhornia paniculata* plants used in this study**

| Individual or population | Location | Population size | Note |
|---|---|---|---|
| Maternal parent of the *SsMm* individual | Yara, Granma, Cuba | | |
| Paternal parent of the *SsMm* individual | Igaci, Alagoas, Brazil | | |
| The *ssmm* individual | San Mateo del Mar, Oaxaca, Mexico | | |
| Pop 1 | Quixadá-2, Ceará, Brazil. Roadside small lake | 200 | Bulk from 30 plants |
| Pop 2 | Quixadá-1, Ceará, Brazil. Roadside marsh | 250 | Bulk from 30 plants |

**Table S5 | Statistics for the two genome assemblies of *Eichhornia paniculata***

| **Genotype of the individual** | *SsMm* | *ssmm* |
|---|---|---|
| **Sequencing technology and depth** | PacBio CLR, 77 Gb (63X) | PacBio CCS (HiFi), 60 Gb (49X) |
| **Assemblers** | Canu + purge_haplotigs | hifiasm + purge_haplotigs |

| | | |
|---|---|---|
| **Contig N50** | 3.81 Mb | 96.8 Mb |
| **Proximity ligation method** | Hi-C + falcon_phase + Proximo | Omni-C + HiRise |
| **Number of scaffolds (pseudomolecules)** | 8 | 8 |
| **Unscaffolded contigs** | 417 | 11 |
| **Total length** | 1,273,901,216 bp | 1,189,336,515 bp |
| **BUSCO scores (dataset: embryophyta_odb10)** | C:98.1%[S:70.4%, D:27.7%], F:1.0%, M:0.9%, n:1614 | C:98.6%[S:85.8%, D:12.8%], F:0.4%, M:1.0%, n:1614 |

**Table S6 | Number of the individuals *N*(indiv) and floral buds *N*(bud) used for each RNA-Seq sample**

| Morph | Tissue | Development stage | Biological replicate 1 (from Pop 1) | | Biological replicate 2 (from Pop 1) | | Biological replicate 3 (from Pop 1) | | Biological replicate 4 (from Pop 2) | |
|---|---|---|---|---|---|---|---|---|---|---|
| | | | *N*(indiv) | *N*(bud) | *N*(indiv) | *N*(bud) | *N*(indiv) | *N*(bud) | *N*(indiv) | *N*(bud) |
| L | Style | Early | 14 | 14 | 14 | 14 | 14 | 14 | 17 | 17 |
| | | Late | 14 | 14 | 14 | 14 | 13 | 14 | 17 | 17 |
| | Stamen | Early | 14 | 14 | 14 | 14 | 14 | 14 | 17 | 17 |
| | | Late | 14 | 14 | 14 | 14 | 13 | 14 | 17 | 17 |
| M | Style | Early | 13 | 13 | 13 | 13 | 9 | 9 | 11 | 22 |
| | | Late | 13 | 13 | 13 | 13 | 13 | 13 | 11 | 11 |
| | Stamen | Early | 13 | 13 | 13 | 13 | 9 | 9 | 11 | 11 |
| | | Late | 13 | 13 | 13 | 13 | 13 | 13 | 11 | 11 |

| | | | | | | | | | | |
|---|---|---|---|---|---|---|---|---|---|---|
| S | Style | Early | 7 | 21 | 7 | 14 | 7 | 21 | 5 | 20 |
| | | Late | 7 | 21 | 7 | 14 | 7 | 13 | 7 | 13 |
| | Stamen | Early | 7 | 7 | 7 | 7 | 7 | 7 | 5 | 5 |
| | | Late | 7 | 7 | 7 | 7 | 7 | 14 | 7 | 13 |

*Early: bud size = 3-5 mm. Late: bud size = 5-8 mm
